# Role Of The C-C Motif Chemokine Ligand 5 (CCL5) And Its Receptor, C-C Motif Chemokine Receptor 5 (CCR5) In The Genesis Of Aldosterone-induced Hypertension, Vascular Dysfunction, And End-organ Damage

**DOI:** 10.1101/2023.09.22.558020

**Authors:** Rafael M. Costa, Débora M. Cerqueira, Ariane Bruder-Nascimento, Juliano V. Alves, Wanessa A.C. Awata, Shubhnita Singh, Alexander Kufner, Eugenia Cifuentes-Pagano, Patrick J. Pagano, Jacqueline Ho, Thiago Bruder-Nascimento

## Abstract

**Background:** Aldosterone, a mineralocorticoid steroid hormone, has been described to initiate cardiovascular diseases by triggering exacerbated sterile vascular inflammation. The functions of C-C Motif Chemokine Ligand 5 (CCL5) and its receptor, C-C Motif Chemokine Receptor 5 (CCR5), are well known in infectious diseases, but their roles in the genesis of aldosterone-induced vascular injury and hypertension are unknown.

**Methods:** We analyzed the vascular profile, blood pressure, and renal damage in wild-type (CCR5^+/+^) and CCR5 knockout (CCR5^−/−^) mice treated with aldosterone (600 µg/kg/day for 14 days) while receiving 1% saline to drink.

**Results:** Here, we show that CCR5 plays a central role in aldosterone-induced vascular injury, hypertension, and renal damage. Long-term infusion of aldosterone in CCR5^+/+^ mice resulted in exaggerated CCL5 circulating levels and vascular CCR5 expression. Aldosterone treatment also triggered vascular injury, characterized by endothelial dysfunction and inflammation, hypertension, and renal damage. Mice lacking CCR5 were protected from aldosterone-induced vascular damage, hypertension, and renal injury. Mechanistically, we demonstrated that CCL5 increased NADPH oxidase 1 (Nox1) expression, reactive oxygen species (ROS) formation, NFκB activation, and inflammation and reduced nitric oxide production in isolated endothelial cells. These effects were abolished by antagonizing CCR5 with Maraviroc. Finally, aortae incubated with CCL5 displayed severe endothelial dysfunction, which is prevented by blocking Nox1, NFκB, or with Maraviroc treatment.

**Conclusions:** Our data demonstrate that CCL5/CCR5, through activation of NFkB and Nox1, is critically involved in aldosterone-induced vascular and renal damage and hypertension. Our data place CCL5 and CCR5 as potential targets for therapeutic interventions in conditions with aldosterone excess.

## Introduction

Aldosterone is a mineralocorticoid steroid hormone, and it is principally generated by the adrenal glomerulosa^1^, although other cell types, including adipocytes^2, 3^, can be an alternative source of aldosterone. The primary function of aldosterone, via mineralocorticoid receptor (MR), is to regulate salt and water homeostasis in the body by acting on the distal tubule and collecting duct of nephrons in the kidney, which in turn leads to sodium and water reabsorption and potassium excretion^1, 4, 5^. However, MR are expressed in different cell types including vascular smooth muscle cells (VSMC)^6^, endothelial cells^9^, adipocytes that form perivascular adipose tissue^7^, and immune cells^8, 9^, where aldosterone can exert its deleterious effects including inflammation, oxidative stress, accelerated fibrosis, and proliferation^4, 5, 8, 10–13^. In this context, hyperaldosteronism and hyperactivation of MR signaling are key features in the pathogenesis of diabetes^2, 3, 10^, obesity^2, 3, 10^, atherosclerosis^14^, stroke^13^, and hypertension^15^. Although such evidence is well-established, the cellular and molecular mechanisms by which aldosterone induces cardiovascular injury are not fully elucidated.

Chemokines (or chemotactic cytokines) are a large family of small proteins that act through cell surface G protein-coupled chemokine receptors^16^. They are best known for their ability to stimulate the migration of cells, most notably immune cells into the injured area^17^. Conversely, chemokines can induce cellular changes independent of immune cell recruitment by signaling directly through their receptors in VSMC and endothelial cells^17–20^.

Chemokine (C-C motif) ligand 5 (CCL5) (or regulated on activation normal T cell expressed and secreted - RANTES) is one of the most important chemokines secreted by T cells, macrophages, activated platelets, endothelial cells, and VSMC^17, 20, 21^. While CCL5 can bind to CCR1, CCR3, CCR4 and CCR5, it has the highest affinity to CCR5, which is expressed in endothelial cells as well^17, 18, 20, 21^. High levels of CCL5 have been identified in hyperlipidemia^22^, atherosclerosis^23, 24^, and hypertension^25^. *In vivo* and *in vitro* studies have revealed that lack or blockage of CCL5 or CCR5 significantly attenuates atherosclerosis^26–28^, neointima formation^29^, atherogenic phenotype switching in obesity^30^, vascular inflammation in acquired lipodystrophy^31^, and inflammation of perivascular adipose tissue in hypertension^25^. We previously reported that CCL5 via CCR5 induces VSMC proliferation and migration in a NADPH oxidase 1 (Nox1)-derived reactive oxygen species (ROS) dependent-manner, whereas blockage of CCR5 *in vivo* attenuates vascular hypertrophy^19^. However, whether the effects of aldosterone-induced vascular injury and hypertension are mediated by exaggerated activity or increased levels of CCL5 and CCR5 is not known.

Herein, we addressed a new mechanism of aldosterone-induced endothelial dysfunction, hypertension, and renal damage. We found that CCL5 and CCR5 are connected to hyperaldosteronism-associated cardiovascular risk. Importantly, high levels of CCL5 in aldosterone-treated mice induces endothelial dysfunction and inflammation via nuclear factor kappa B (NFκB) and Nox1-derived ROS, whereas deficiency in CCR5 protects against aldosterone-induced vascular injury, hypertension, and end-organ damage. Therefore, we place CCL5 and CCR5 as a major trigger of cardiovascular disease in conditions associated with aldosterone excess such as primary aldosteronism, obesity, and hypertension.

## Methods

### Mice

Ten- to twelve-week-old male CCR5^+/+^ (C57BL6/J) and CCR5^−/−^ (Ccr5tm1Kuz/J Jax #005427) mice were used. All mice were fed with standard mouse chow and tap water was provided *ad libitum*. Mice were housed in an American Association of Laboratory Animal Care-approved animal care facility in Rangos Research Building at Children’s Hospital of Pittsburgh (CHP) of University of Pittsburgh. The Institutional Animal Care and Use Committee approved all protocols (IACUC protocols #19065333 and #22061179). All experiments were performed in Rangos Research Building at CHP and were in accordance with the Guide Laboratory Animals for The Care and Use of Laboratory Animals.

### Aldosterone treatment

CCR5^+/+^ and CCR5^−/−^ mice were infused with vehicle (saline) or aldosterone (600 μg/kg per day) for 14 days with ALZET osmotic minipumps (Durect, Cupertino, CA) while receiving 1% saline in the drinking water as described previously^8^.

### Circulating chemokine profile

Circulating chemokines levels were analyzed via the Proteome Profiler Mouse Cytokine Array from R&D System. Serum was pooled from six CCR5^+/+^ mice treated with vehicle or aldosterone. Data were presented as fold changes by a Heat Map graph.

### Circulating CCL5 levels

Circulating CCL5 levels were measured in serum from CCR5^+/+^ mice treated with vehicle or aldosterone via Enzyme-Linked Immunosorbent Assay (ELISA) (R&D System).

### Vascular function

As described previously^32^ ^33^, thoracic aortae from CCR5^+/+^ and CCR5^−/−^ mice treated with vehicle or aldosterone were dissected from connective tissues, separated into rings (2 mm), and mounted in a wire myograph (Danysh MyoTechnology) for isometric tension recordings with PowerLab software (AD Instruments). Rings were placed in tissue baths containing warmed (37 °C), aerated (95% O_2_, 5% CO_2_) Krebs Henseleit Solution (in mM: 130 NaCl, 4.7 KCl, 1.17 MgSO_4_, 0.03 EDTA, 1.6 CaCl_2_, 14.9 NaHCO_3_, 1.18 KH_2_PO_4_, and 5.5 glucose). After 30 min of stabilization, KCl (120 mM) was used to test arterial viability. Concentration-effect curves for phenylephrine (PE, α-1 adrenergic receptor-dependent vasoconstrictor), acetylcholine (ACh, endothelium-dependent vasodilator), and sodium nitroprusside (SNP, endothelium-independent vasodilator) were performed. Some experiments were performed in the presence of Nox1Ads (10 µM, specific Nox1 inhibitor) to analyze the involvement of Nox1 on aldosterone-induced vascular dysfunction.

#### Ex vivo protocol for isolated aortae incubation

Aortae were harvested from CCR5^+/+^ mice, separated in 2 mm rings, and incubated with CCL5 (100 ng/mL) for 24 hours. Some aortic rings were treated with Maraviroc (40 µM, specific CCR5 antagonist), Nox1Ads (10 µM), or BMS-345541 (5 µM, NFκB signaling inhibitor). Then, experiments of endothelial function were performed.

### Blood pressure analysis

CCR5^+/+^ and CCR5^−/−^ mice were instrumented with telemetry transmitters to record arterial pressure and heart rate (HD-X10, Data Sciences International). Transmitters were implanted as described previously^34–36^. After 7 days of recovery from surgery, necessary for the mice to gain their initial body weight, data were recorded for 4 days as baseline. Then, aldosterone was infused via osmotic mini-pump for 14 days as described^8^, and blood pressure [systolic blood pressure (SBP), diastolic blood pressure (DBP) and mean arterial pressure (MAP)] values were obtained for 2-3 hours per day between 12pm-5pm.

### Immunohistochemical staining for renal injury

As described previously^37^, kidneys were fixed in 4% paraformaldehyde overnight, embedded in paraffin and sectioned at 4 μm. After deparaffinization, rehydration, and permeabilization in PBS containing 0.1% Tween-20 (PBS-T), antigen retrieval was performed by boiling the slides for 30 min in either 10 mM sodium citrate pH 6.0 buffer or Trilogy (Cell Marque). Sections were then blocked in 3% BSA before and incubated overnight with the appropriate primary antibody at 4°C. Then, sections were washed with PBS-T, incubated with secondary antibody, washed again with PBS-T and mounted in Fluoro Gel with DABCO™ (Electron Microscopy Science) before visualizing with either a Leica DM2500 microscope equipped with a Leica DFC 7000T camera and a LAS X software or with a Zeiss 710 confocal microscope with Zen software (Zeiss). The antibodies used for immunostaining are shown in Supplementary Table 1. Primary antibodies were visualized by staining with fluorescence-conjugated antibodies. Nuclei were counterstained with DAPI. Proximal tubules were visualized with fluorescein-labeled Lotus tetragonolobus lectin (LTA).

### Proteinuria

Urine was collected during the tissue harvesting, stored at −80°C, and proteinuria was analyzed as previously described^38^. 15 µL of urine samples were separated on 10% SDS-PAGE gels followed by staining with Coomassie Brilliant Blue (Sigma-Aldrich). Albumin (15 µg) was used as a control.

### Endothelial cell culture

Mouse Mesenteric Endothelial Cells (MEC) were purchased from Cell Biologics. Cells were maintained in Complete Mouse Endothelial Cell Medium (Cell Biologics) containing Endothelial Cell Medium Supplement Kit (Cell Biologics). Cells were used between passage 4-8.

#### Protocol of endothelial cells stimulation

MEC were treated with CCL5 (100 ng/mL) for 24 hours in the presence of vehicle or Maraviroc (40 µM), Nox1Ads (10 µM), or BMS-345541 (5 µM). MEC were also treated with aldosterone (0.1 µM) with or without eplerenone (1 µM). MEC were incubated for 30 min with Inhibitors and antagonists prior to CCL5 or aldosterone treatments.

### Macrophage adhesion assay

Endothelial macrophage adhesion was determined according to our previously described methods^11^ ^39^. Briefly, MEC were cultured to confluence in 6-well plates and treated with CCL5 (100 ng/mL) for 24 hours in the presence of vehicle or Maraviroc (40 µM). Non-stimulated MEC served as controls. Macrophage cells (RAW 264.7, ATCC) were cultured in Dulbecco’s Modified Eagle’s Medium (DMEM) contained 10% of Fetal Bovine Serum (FBS). For cell fluorescent labeling, macrophages (10^5^ cells/mL) were suspended in 1% bovine serum albumin (BSA)-supplemented phosphate buffered saline containing 1 μM calcein-AM (Invitrogen) and incubated for 20 minutes at 37 °C. Labeled macrophages were washed twice with phosphate-buffered saline and suspended in Hanks’ buffered salt solution. Fluorescence labeled cells (10^5^ cells/well) were then added to both non-stimulated and stimulated MEC layers and were allowed to adhere for 30 minutes at 37 °C in 5% CO_2_. After incubation, non-adhered cells were removed by gently washing with pre-warmed Hanks’ buffered salt solution. The number of adherent cells was determined by images or fluorescence intensity via fluorescence microscopy (Leica Microsystems) and fluorimeter (SpectraMax i3x Multi-Mode Microplate Reader), respectively. For the fluorescence intensity, cells were lysed with 0.1 M NaOH and transferred to a 96 well plate, then fluorescence intensity was measured.

### ROS measurement

Aortae and MEC were collected in lysis buffer (Hank’s Balanced Salt Solution with Complete Mini protease inhibitor and PhosSTOP phosphatase inhibitor) and were lysed by five freeze/thaw cyclesand passed through a 30-gauge needle five times to disrupt cells to measured ROS as described before^40^. The cell lysates were centrifuged at 1000 g for 5 min at 4 °C to remove unbroken cells, nuclei, and debris. Throughout all procedures, extreme care was taken to maintain the lysate at a temperature close to 0 °C. Lysates of aortae and MEC were resuspended in Amplex Red assay mixture (25 mM HEPES, pH 7.4, containing 120 mM NaCl, 3 mM KCl, 1 mM MgCl2, 0.1 mM Amplex red (Invitrogen), and 0.35 U/ml horseradish peroxidase (HRP) in the presence and absence of catalase (300 U/ml). The reaction was initiated by the addition of 36 µmol/l NADPH (MP Biomedicals). Florescence measurements were made using a Biotek Synergy 4 hybrid multimode microplate reader with a 530/25-exitation and a 590/35-emission filter. The reaction was monitored at 25 °C for 1 hour.

### Nitric oxide measurement

Nitric oxide production was measured by 4,5-Diaminofluorescein diacetate (DAF-2 DA) probe. Briefly, MEC were treated with CCL5 (100 ng/mL) for 24 hours in the presence of vehicle or Maraviroc (40 µM). Then, media were replaced, and bradykinin (1 µM)^41^ was used to stimulate nitric oxide production for 30 minutes. After the treatments, cells were washed with PBS and stained with DAF-2 DA (5 µM) for 30 minutes before the analysis. Bradykinin (5 µM) was used as positive control. Fluorescence intensity was measured in a fluorimeter (SpectraMax i3x Multi-Mode Microplate Reader) (Emission. 538 nm/Excitation. 485 nm).

### Western Blot

Aortic protein was extracted using radioimmunoprecipitation assay buffer (RIPA) buffer (30 mM HEPES, pH 7.4,150 mM NaCl, 1% Nonidet P-40, 0.5% sodium deoxycholate, 0.1% sodium dodecyl sulfate, 5 mM EDTA, 1 mM NaV0_4_, 50 mM NaF, 1 mM PMSF, 10% pepstatin A, 10 μg/mL leupeptin, and 10 μg/mL aprotinin). Total protein extracts were centrifuged at 15,000 rpm/10 min and the pellet was discarded. Proteins from homogenates of aortae (25 μg) were used. MEC samples were directly homogenized using 2x Laemmli Sample Buffer and supplemented with 2-mercaptoethanol (β-mercaptoethanol) (BioRad Hercules). Proteins were separated by electrophoresis on a polyacrylamide gradient gel (BioRad Hercules) and transferred to Immobilon-P poly (vinylidene fluoride) membranes. Non-specific binding sites were blocked with 5% skim milk or 1% bovine serum albumin (BSA) in tris-buffered saline solution with tween for 1h at 24 °C. Membranes were then incubated with specific antibodies overnight at 4 °C as described in Supplementary Table 2. After incubation with secondary antibodies, the enhanced chemiluminescence luminol reagent (SuperSignal™ West Femto Maximum Sensitivity Substrate, Thermo Fisher) was used for antibody detection.

### Real-Time Polymerase Chain Reaction (RT-PCR)

mRNA from aortae and MEC were extracted using RNeasy Mini Kit (Quiagen). Complementary DNA (cDNA) was generated by reverse transcription polymerase chain reaction (RT-PCR) with SuperScript III (Thermo Fisher). Reverse transcription was performed at 58 °C for 50 min; the enzyme was heat inactivated at 85 °C for 5 min, and real-time quantitative RT-PCR was performed with the PowerTrack™ SYBR Green Master Mix (Thermo Fisher). Sequences of genes as listed in Supplementary Table 3. Experiments were performed in a QuantStudio™ 5 Real-Time PCR System, 384-well (Thermo Fisher). Data were quantified by 2ΔΔ Ct and are presented by fold changes indicative of either upregulation or downregulation.

### Statistical analysis

For comparisons of multiple groups, one-way or two-way analysis of variance (ANOVA), followed by the Tukey post-test was used. Differences between the two groups were determined using Student’s t-test. The vascular relaxation response is expressed as a percentage of relaxation based on the phenylephrine maximal response (1uM), whereas contractile response is presented as millinewton (mN). Maximal response (Emax) and negative logarithm of EC50 (*p*D_2_) were determined. Analyses were performed using Prism 10.0 software (GraphPad). A difference was considered statistically significant when p ≤ 0.05.

## RESULTS

### Circulating CCL5 levels and vascular CCR5 expression are increased in aldosterone-treated mice and may play important role on aldosterone-induced endothelial dysfunction

We firstly observed that CCR5^+/+^ mice treated with aldosterone displayed changes in circulating cytokines and chemokines (Fig. 1A). However, CCL5, which is considered the main CCR5 ligand, was the most elevated in the proteome cytokine array. In addition, we also used ELISA to confirm the high CCL5 circulating levels post-aldosterone treatment (Fig. 1B).

**Figure 1.**
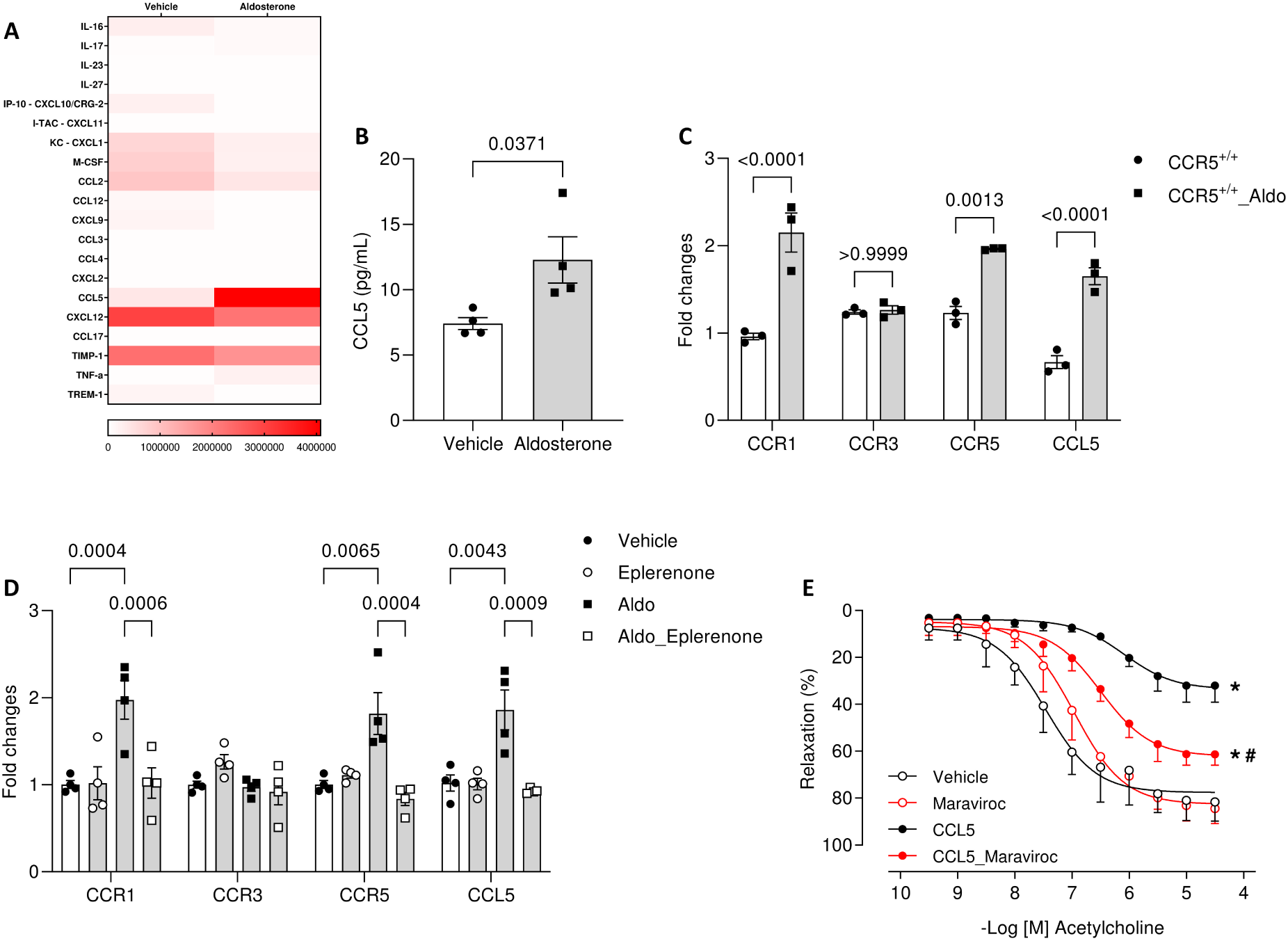
Aldosterone-induced CCL5 high levels promotes endothelial dysfunction. Inflammatory profile in plasma, measured by proteome profiler mouse cytokine array and presented as heat map (A); CCL5 levels, measured by ELISA (B); and chemokine receptors expression, measured by RT-PCR (C), from CCR5^+/+^ mice treated with vehicle or aldosterone (600 µg/kg/day for 14 days). Chemokine receptors expression, measured by RT-PCR, in endothelial cells (MEC) treated with vehicle or aldosterone (0.1µM) in the presence of Eplerenone (1µM) (D). Concentration-effect curves to acetylcholine in aortae from CCR5^+/+^ mice incubated with vehicle or CCL5 (100ng/ml, 24h) in the presence of Maraviroc (40µM) (E). Values represent means ± SEM (n= 3-7). Student t test or ANOVA test. *p<0.05 *vs*. Vehicle; #p<0.05 *vs.* CCL5.

In aortae from aldosterone-treated mice, we found elevated CCR1, CCR5, and CCL5 gene expression, with no difference for CCR3 (Fig. 1C). To understand whether aldosterone regulates chemokine receptors expression, we treated endothelial cells with aldosterone and observed a significant increase in CCR1, CCR5, and CCL5 levels, which was dependent on MR activation, since eplerenone blunted the aldosterone effects (Fig. 1D). These data indicate that aldosterone increases CCL5 production and CCR5 expression in endothelial cells in a MR dependent manner.

We recently demonstrated that CCL5 induces vascular inflammation and proliferation dependent on CCR5. Although CCL5 has a large affinity by CCR5, other chemokine receptors can recognize CCL5, such as CCR1 and CCR3. Therefore, we investigated whether CCL5 is leading to endothelial dysfunction via CCR5. By incubating aortae from CCR5^+/+^ mice with CCL5, with or without a specific CCR5 antagonist (Maraviroc), and performing studies of endothelial function by myography, we observed that CCL5 triggered severe endothelial dysfunction in aortae, which was partially protected by antagonizing CCR5 (Fig. 1E). Furthermore, CCL5 incubation for 24h elevated CCR1 and CCL5 expressions in isolated aortae (Supplementary Fig. 1A). CCL5 triggered vascular hypercontractile to phenylephrine in aortae, which was protected by antagonizing CCR5 (Supplementary Fig. 1B). No difference was observed for SNP (endothelium-independent vasodilator) (Supplementary Fig. 1C).

### CCR5 deficiency protects from aldosterone-induced endothelial dysfunction and vascular inflammation

Since we found that CCL5 and CCR5 are elevated in aldosterone-treated mice and CCL5 induces endothelial dysfunction in a CCR5-dependent manner, we treated CCR5^+/+^ and CCR5^−/−^ mice with aldosterone to analyze whether lack in CCR5 might confer protection against vascular injury. Interestingly, we found that aldosterone promoted severe vascular dysfunction characterized by an impaired endothelium-dependent relaxation to ACh (Fig. 2A) and hypercontractility to phenylephrine (Supplementary Fig. 2A); these changes were prevented by CCR5 deficiency. No difference was observed for SNP (Fig. 2B).

**Figure 2.**
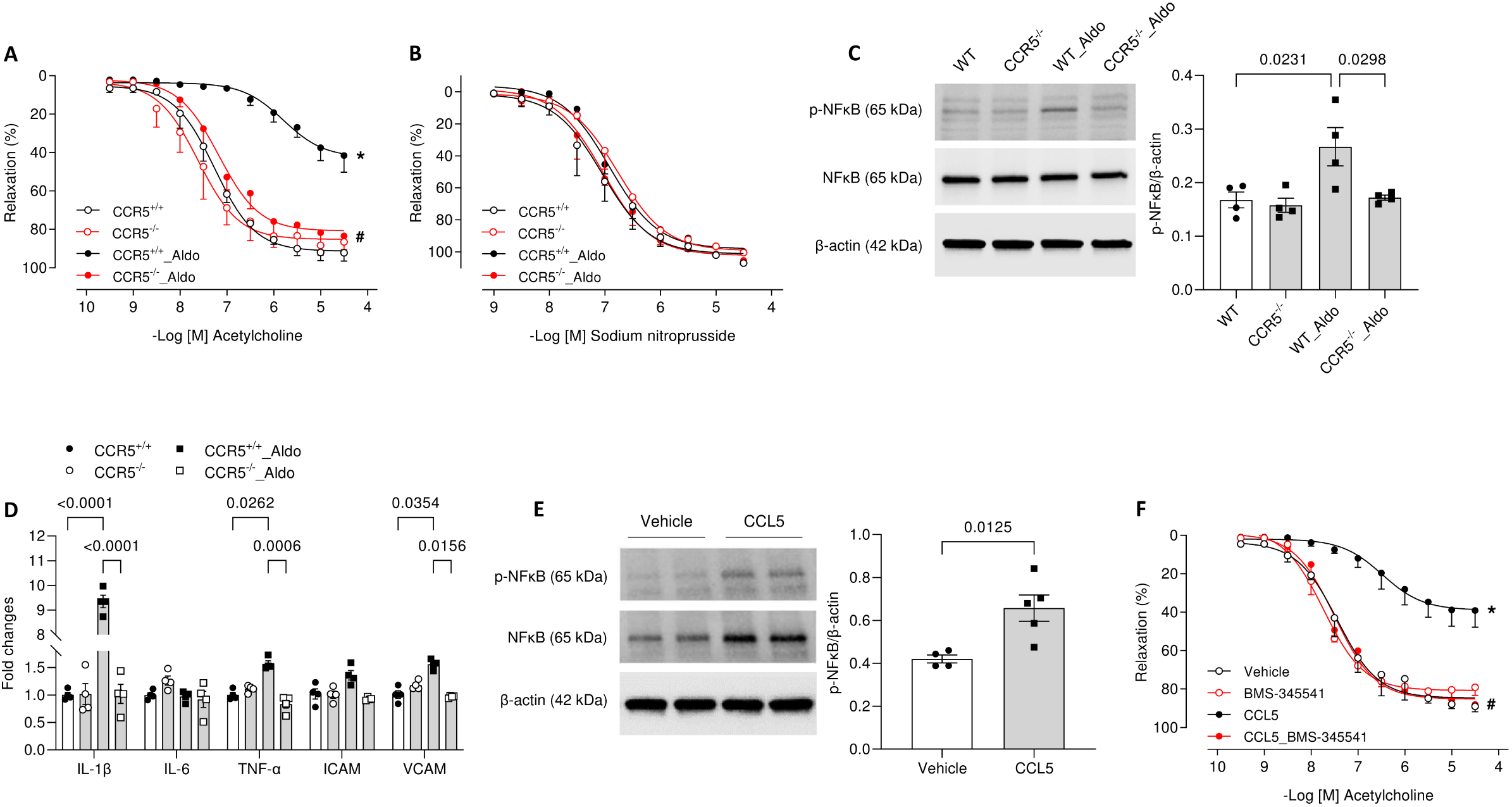
CCR5 deficiency prevents endothelial dysfunction and vascular inflammation induced by aldosterone. Concentration-effect curves to acetylcholine (A) and sodium nitroprusside (B); phosphorylated (p65 subunit) and total NFκB expression, analyzed by western blot to (C); inflammatory markers, measured by RT-PCR (D), in aortae from CCR5^+/+^ and CCR5^−/−^ mice treated with vehicle or aldosterone (600 µg/kg/day + saline for 14 days). Phosphorylated (p65 subunit) and total NFκB expression analyzed by western blot in aortae from CCR5^+/+^ mice incubated with vehicle or CCL5 (100ng/ml, 24h) (E) and concentration-effect curves to acetylcholine (F) in aortae from CCR5^+/+^ mice incubated with vehicle or CCL5 (100ng/ml, 24h) in the presence of BMS-345541 (5µM). Values represent means ± SEM (n= 4-7). Student t test or ANOVA test. *p<0.05 *vs.* CCR5^+/+^ or Vehicle; #p<0.05 *vs.* CCR5^+/+^_Aldo or CCL5.

We found that aldosterone treatment augmented NFkB phosphorylation in aortae from CCR5^+/+^ mice, but not in CCR5^−/−^ mice (Fig. 2C). These effects were accompanied by increased expression of inflammatory genes (IL-1β, TNF-α, and VCAM expression) (Fig. 2D), which was not detected in CCR5^−/−^ mice. *Ex vivo* experiments in aortae revealed that CCL5 incubation for 24h increased NFkB activation (Fig. 2E), whereas NFkB inhibition prevented CCL5-induced endothelial dysfunction (Fig. 2F). Furthermore, CCL5 incubation for 24h increased expression of inflammatory genes (IL-1β, TNF-α, ICAM and VCAM expression) in isolated aortae (Supplementary Fig. 2B).

To confirm that CCL5/CCR5 leads to endothelial inflammation, we interrogated whether CCL5 triggers endothelial inflammation by treating MEC with CCL5 in the presence or absence of Maraviroc and analyzing inflammatory markers and RAW264.7 adhesion. CCL5 augmented IL1β, TNF-α, ICAM and VCAM gene expression and increased RAW264.7 adhesion in MEC at 24 hr, whereas Maraviroc prevented these effects (Fig. 3A and B).

**Figure 3.**
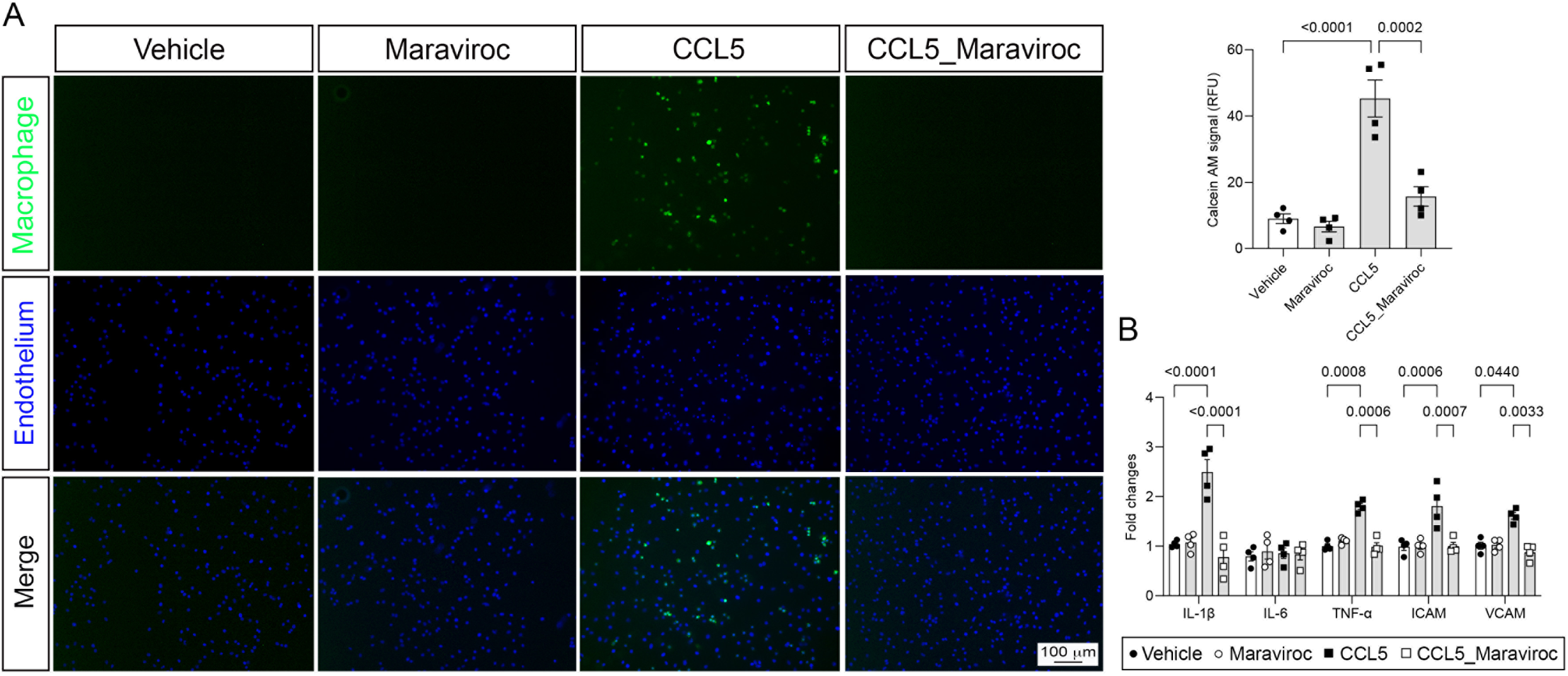
CCL5 via CCR5 induces endothelial cell activation and immune cell adhesion. Photomicrography and fluorescence intensity depicting labeled macrophages (calcein-AM probe, green) and endothelial cells (MEC, DAPI, blue). MEC were treated with vehicle, Maraviroc (40µM, 30 minutes prior CCL5 incubation), CCL5 (100ng/ml, 24h) or CCL5 plus Maraviroc-stimulated MEC. Scale bar = 100µm (A). Inflammatory markers, measured by RT-PCR, in MEC treated with vehicle or CCL5 (100ng/ml) in the presence of Maraviroc (40µM) (B). Values represent means ± SEM (n= 4). ANOVA test.

### CCR5 deficiency prevents aldosterone-induced hypertension and kidney damage

Endothelial dysfunction is a key feature in the genesis and progression of hypertension and end-organ damage. Therefore, we investigated whether CCR5 deficiency could protect from aldosterone-induced hypertension and renal injury. Aldosterone induced hypertension is characterized by increased SBP, DBP, and MAP. Interestingly, CCR5^−/−^ mice were protected against aldosterone-induced hypertension (Fig. 4A-C).

**Figure 4.**
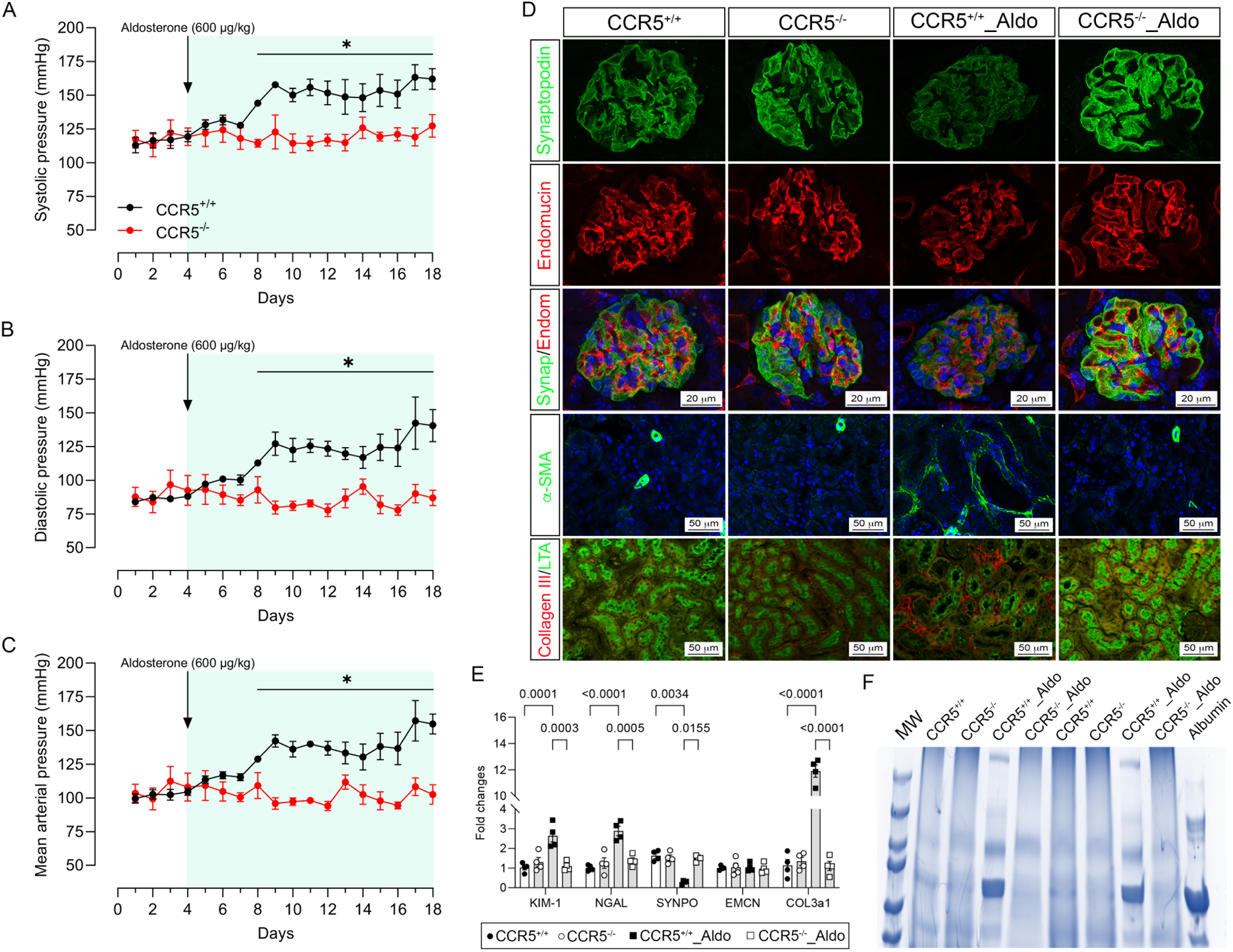
Aldosterone via CCR5 promotes increased blood pressure and kidney injury. Systolic blood pressure (A), diastolic blood pressure (B), and mean arterial pressure (C), measured via radiotelemetry, in CCR5^+/+^ and CCR5^−/−^ mice treated with vehicle or aldosterone (600 µg/kg/day and saline for 14 days). Immunofluorescence for podocyte marker synaptopodin (Synap; green), endothelial marker endomucin (Endom; red), and nuclei (DAPI, blue) represented by letters A to D, α-smooth muscle actin (α-SMA; green) and nuclei (DAPI, blue) represented by letters E to H, fibrosis marker and proximal tubules were visualized with fluorescein-labeled collagen III (red) and Lotus tetragonolobus lectin (LTA; green), which are represented by letters I to L. Images were obtained from kidney sections of CCR5^+/+^ and CCR5^−/−^ mice treated with vehicle or aldosterone. The images shown are representative of three independent experiments. Scale bar = 20 or 50µm (D). Kidney injury markers, measured by RT-PCR in CCR5^+/+^ and CCR5^−/−^ mice treated with vehicle or aldosterone (E), and proteinuria levels in CCR5^+/+^ and CCR5^−/−^ mice treated with vehicle or aldosterone measured by 10% SDS-PAGE gels followed by staining with Coomassie Brilliant. Albumin was used as a control analyzed. Values represent means ± SEM (n= 3-7). Student t test or ANOVA test. *p<0.05 *vs.* CCR5^+/+^.

In kidneys, aldosterone augmented the expression of the renal injury markers, Kidney injury molecule-1 (KIM-1) and Neutrophil gelatinase-associated lipocalin (NGAL) by qRT-PCR (Fig. 4E). Furthermore, immunostaining analyses indicated elevated tubulointerstitial deposition of fibrotic markers, αSMA and collagen III (Fig. 4D). Increased collagen III levels were also confirmed by qRT-PCR (Fig. 4E). Interestingly, aldosterone-treated mice exhibited reduced expression of the podocyte-specific marker synaptopodin (measured by immunofluorescence and RT-PCR) (Fig. 4D and Fig. 4E), and this was accompanied by augmented proteinuria (Fig. 4F). No changes in the expression of the endothelial cell marker, endomucin, were detected (Fig. 4D and Fig. 4E). All these deleterious effects caused by aldosterone (renal damage and proteinuria) were abolished in CCR5^−/−^ mice. Finally, no structural difference was observed in the afferent and efferent glomerular arterioles (Supplementary Fig. 3). These data suggest that CCL5/CCR5 play major role on aldosterone-associated hypertension and renal damage.

### Aldosterone induces endothelial injury and Nox1-derived ROS and impairs nitric oxide formation via CCL5 and CCR5

Aldosterone has been associated with Nox activation in different cell types. Herein, we demonstrated that aldosterone treatment increases aortic Nox1 (but not Nox2 and Nox4) (Fig. 5A) and induces ROS production (Fig. 5B) in CCR5^+/+^ mice. CCR5^−/−^ mice were protected from these effects. To understand whether aldosterone is leading to endothelial dysfunction via Nox1 activation, we incubated arteries prior to ACh curves with a selective Nox1 inhibitor, NoxA1ds^40^, and observed that Nox1 inhibition blunted the endothelial dysfunction in CCR5^+/+^ mice (Fig.5C).

**Figure 5.**
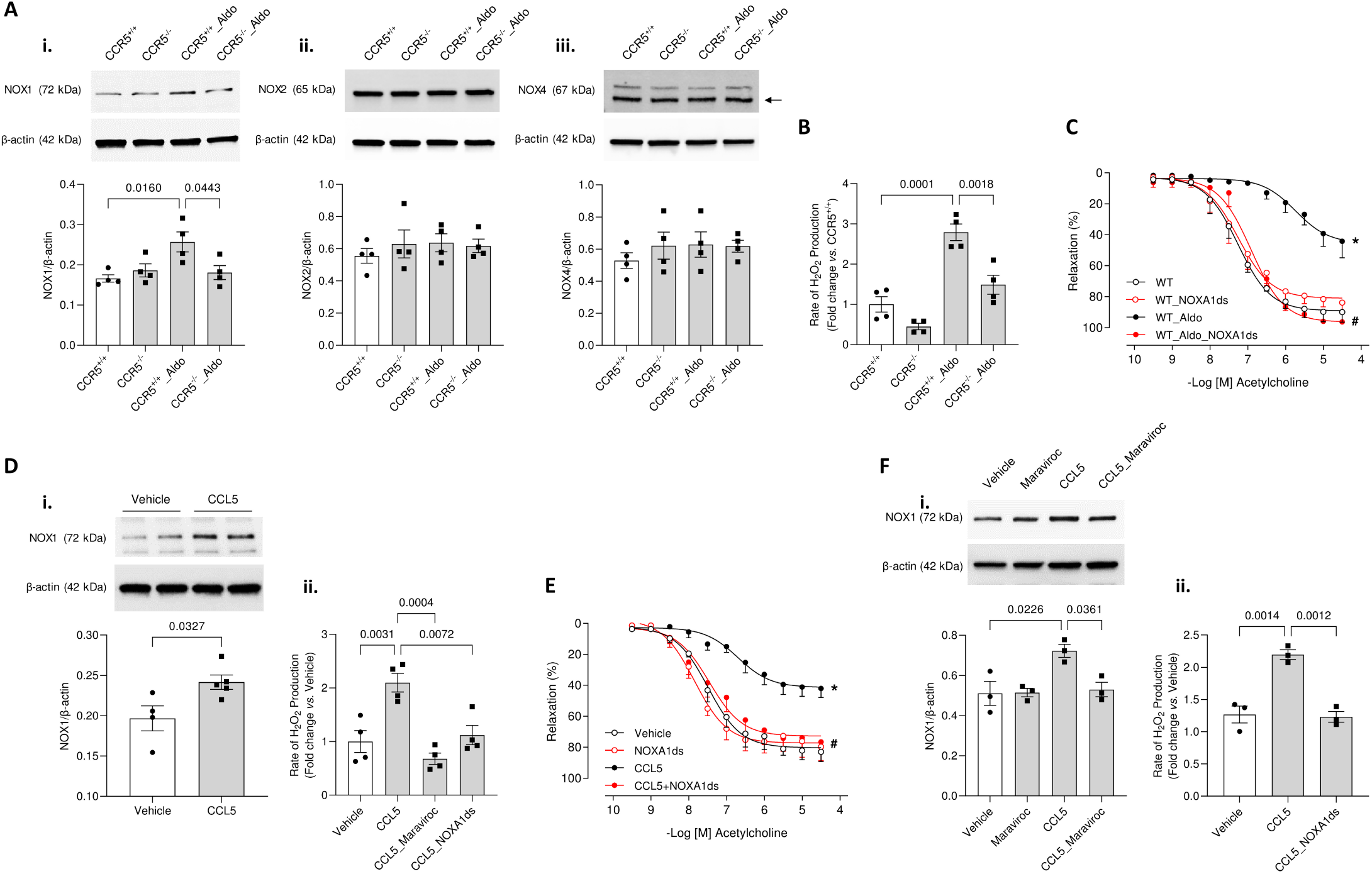
Aldosterone, via CCR5, promotes endothelial dysfunction by NOX1-dependent mechanisms. Representative western blot (A) to NOX1 (i), NOX2 (ii) and NOX4 (iii) expression; ROS generation, measured Amplex red assay (B); and concentration-effect curves to acetylcholine, in the presence of NOXA1ds (10µM) (C), in aortae from CCR5+/+ and CCR5-/- mice treated with vehicle or aldosterone (600 µg/kg/day for 14 days). Representative western blot to NOX1 (i); ROS generation, measured by Amplex red assay (ii) (D); and concentration-effect curves to acetylcholine, in the presence of NOXA1ds (10µM) (E), in aortae from CCR5+/+ mice incubated with vehicle or CCL5 (100ng/ml). Representative western blot to NOX1 (i) and ROS generation, measured by Amplex red (ii), in endothelial cells (MEC) treated with vehicle or CCL5 (100ng/ml) in the presence of Maraviroc (40µM) and NOXA1ds (10µM) (F). Values represent means ± SEM (n= 4-7). ANOVA test. *p<0.05 *vs.* CCR5^+/+^ or Vehicle; #p<0.05 *vs.* CCR5^+/+^_Aldo or CCL5.

Next, we assessed whether CCL5 induces endothelial dysfunction via Nox1 activation by multiple approaches. Firstly, we treated isolated aortae from CCR5^+/+^ mice with CCL5 for 24h, and we observed an increase in the Nox1 expression (Fig. 5D), but not Nox2 and Nox4 (Supplementary Fig. 4A-B), and in the ROS generation (Fig. 5D). This increase in ROS generation was abolished in the presence of a selective Nox1 inhibitor, NoxA1ds (Fig. 5D). In addition, in presence of Nox1Ads, we performed studies of endothelial function to examine whether CCL5 induces endothelial dysfunction via Nox1 and found Nox1 inhibition conferred protection against CCL5-induced endothelial dysfunction (Fig. 5E).

Secondly, we treated MEC with CCL5 in presence of Maraviroc to confirm that CCL5 increases endothelial Nox1 and ROS production. We observed that CCL5 treatment for 24h augmented Nox1 expression and induced ROS formation, which were blunted by blocking CCR5 (Maraviroc)or inhibiting NOX1 (NOXA1ds) (Fig. 5F).

Lastly, we treated MEC with CCL5 to analyze whether it can affect nitic oxide formation, as displayed in Supplementary Figures 4C and 4D. CCL5 (100ng/mL, 24h) decreased eNOS phosphorylation at Ser^1177^ and impaired bradykinin-induced nitric oxide formation at 24h, which were prevented by antagonizing CCR5 with Maraviroc.

### Aldosterone induces Nox1 expression and endothelial dysfunction via CCL5/CCR5 and NFkB

The association between aldosterone, Nox1, and NFκB, as well as NFκB modulating Nox1 expression has been described before, but whether such communications are dependent on CCL5 and CCR5 are unknown. *In vitro*, we found that CCL5 induces an increase in Nox1 expression, which is prevented by the presence of the NFκB pathway inhibitor (Fig. 6A). Furthermore, the presence of the NFκB inhibitor blocks ROS generation (Fig. 6B). By a feedback mechanism, CCL5 increases NFκB phosphorylation, which was reverted in the presence of the Nox1 inhibitor (Fig. 6C), indicating that ROS formed by Nox1 increase NFκB activity, while NFκB activity can regulate NOX1 expression and ROS formation. Therefore, we can suggest that CCL5 induces endothelial injury via a continuous and positive feedback between Nox1 and NFκB.

**Figure 6.**
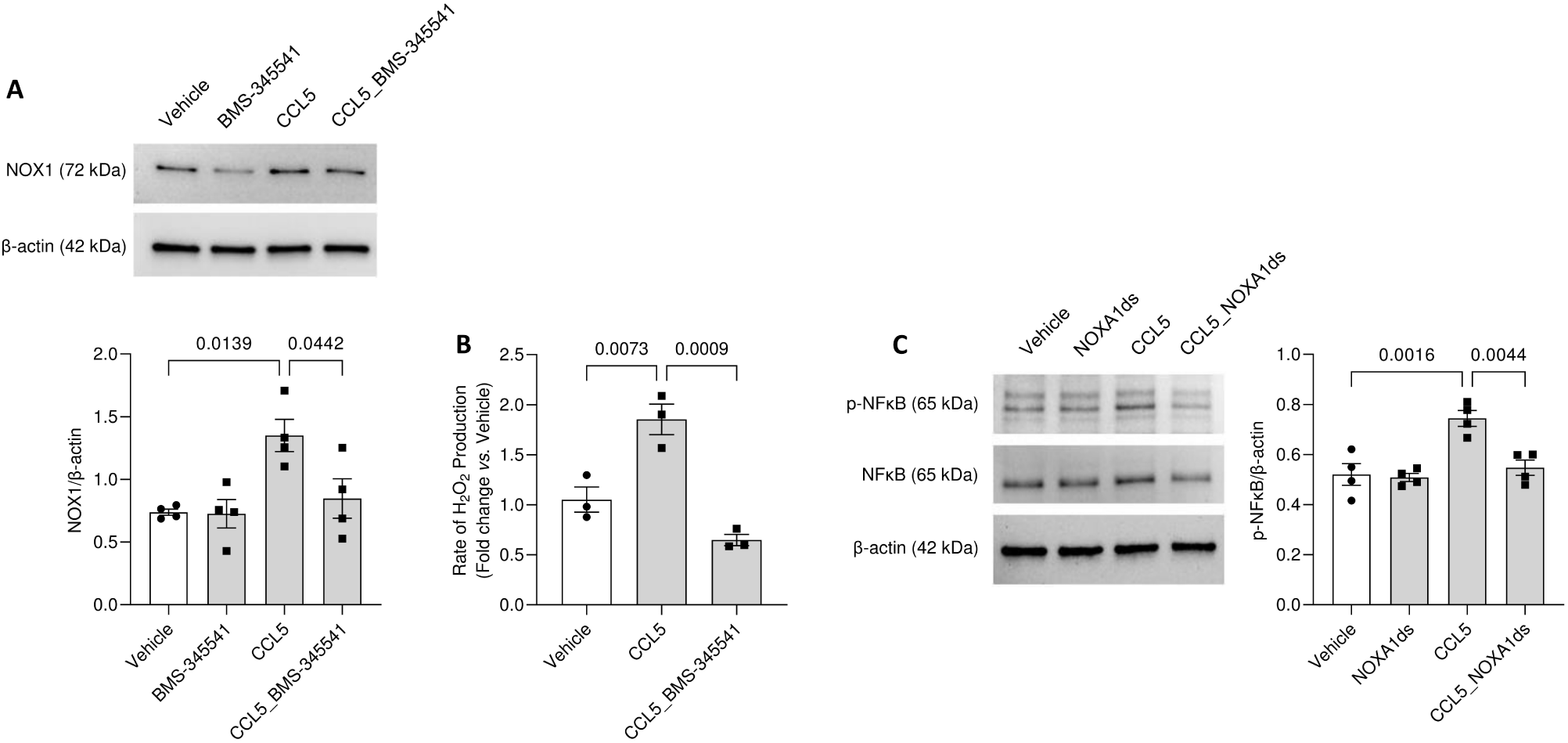
CCL5 induces an increase in NOX1 expression and NFKB activity by a positive feedback mechanism in endothelial cells. Representative western blot to NOX1 expression (A) and ROS generation, measured by Amplex red (B), in endothelial cells (MEC) treated with vehicle or CCL5 (100ng/ml), in the presence of BMS-345541 (5µM). Representative western blot to phosphorylated and total NFκB expression in MEC treated with vehicle or CCL5, in the presence of NOXA1ds (10µM) (C). Values represent means ± SEM (n= 4). ANOVA test.

## Discussion

Cardiovascular diseases are the leading cause of death globally. Aldosterone contributes to the endocrine basis of the development and progression of multiple cardiovascular disease processes, including hypertension, chronic kidney disease, coronary artery disease, and congestive heart failure^4, 42^. The association between aldosterone and genesis of hypertension is particularly strong^3, 4, 15, 43^. Several studies place this mineralocorticoid hormone as a major trigger to the development and severity of hypertension, even in the absence of classically defined primary aldosteronism. For instance, blockage of the aldosterone receptor, MR, decreases blood pressure in rodent models of hypertension^44^, improves cardiovascular and renal outcomes associated with obesity and diabetes^12, 45–49^, restores obesity-associated hypertension^46, 50^. Although the beneficial hemodynamic effects of aldosterone blockage have been described, the molecular and cellular mechanisms remain not fully elucidated.

Aldosterone is a potent inflammatory hormone, and several of its deleterious cardiovascular effects is mediated by its inflammatory capacity, which in turn can lead to dysfunction, fibrosis, and remodeling in the heart, vasculature, and kidney, as well as hypertension^51^. We previously demonstrated that aldosterone induces VSMC inflammation via oxidative stress^11^ and triggers vascular injury and hypertension dependent on NLRP3 inflammasome and IL-1β formation in immune compartment^8^. Others have demonstrated that aldosterone treatment impairs T-regulatory cells (T-Reg) recruitment into injury sites, impairing a tuned innate and adaptive immune response and impacting vascular function and blood pressure control^52^. In atherosclerosis, lack of endothelial MR confers protection against leukocyte-endothelial interactions, plaque inflammation, and expression of adhesion proteins^14^. Therefore, elucidating how aldosterone initiates an inflammatory response is highly important to generate new therapeutic approaches for aldosterone excess-associated cardiovascular disease.

In this study, we are describing a novel mechanism by which aldosterone induces vascular injury, hypertension, and end-organ damage. We are demonstrating that aldosterone (1) increases vascular CCR5 via MR activation and circulating CCL5 levels in mice, (2) induces vascular dysfunction, inflammation, and remodeling in a CCR5-dependent manner, (3) promotes hypertension and renal damage via CCR5, and (4) activates a continuous and positive feedback between NFκB and Nox1-derived ROS in the vasculature via CCL5 and CCR5. These data reveal a remarkable participation of CCL5 and CCR5 in aldosterone-associated cardiovascular damage and hypertension.

Hypertension is associated with immune cell activation and their migration into the kidney, vasculature, heart, and brain; such mechanisms are central for blood pressure regulation and convincing targets for pharmacological intervention^53^. Because chemokines and their receptors (chemokine receptors) help recruit immune cells into the inflamed or injured areas, they are believed to be a leading pathway in the development and progression of hypertension, but whether any interaction between chemokines and their receptors can directly propagate vascular injury, changes in blood pressure, and end-organ damage is unknown.

CCR5 and CCL5 are elevated in cardiovascular events^19, 27–29, 31^ including hypertension^54–56^. Aligned with these previous reports, we observed that aldosterone-treated mice display increased circulating CCL5 and vascular CCR5, which seems to be mediated by MR activation, since aldosterone-induced CCR5 expression in endothelial cells is blunted by eplerenone. Although CCR5 and CCL5 are increased in different models of hypertension, whether they can modulate vascular function, blood pressure control, and renal injury is controversial^25, 55, 57^ or not fully described. Furthermore, it is not clear whether CCR5 can participate in the development and progression of any model of hypertension, or if it is model dependent.

The role of CCR5 and CCL5 on hypertension has been explored before, but not in the aldosterone model. Krebs et al^55^ demonstrated that CCR5 deficient mice are not protected from cardiac and renal injury and hypertension induced by DOCA salt (50mg) + angiotensin II infusion (1500ng/Kg/min), whereas Rudemiller et al^57^ showed that CCL5 knockout mice demonstrate a worse renal outcome post angiotensin II treatment (1000ng/Kg/min) with no changes in blood pressure. Furthermore, Mikolajczyk et al^25^ showed that CCL5 knockout mice or pharmacological intervention with CCL5 blockage (met-RANTES) does not affect blood pressure in angiotensin II-induced hypertension model (490ng/Kg/min), but improves vascular function and decreases vascular ROS, and T-cells infiltration into perivascular adipose tissue. In contrast, we found that CCR5 deficient mice are protected from aldosterone-induced vascular injury, hypertension, and renal damage. This discrepancy may be due to the highly potent model of hypertension, DOCA salt plus a very high dose of angiotensin II treatment used by Krebs et al^55^. On the other hand, Rudemiller et al^57^ or Mikolajczyk et al^25^ used CCL5 deficient mice, and not CCR5. Other ligands than CCL5 can activate CCR5 including CCL3 and CCL4^16^, thus these intriguing studies do not rule out the effect of other chemokines activating CCR5. Finally, none of these studies used an aldosteronism model to induce hypertension and injury, so involvement of CCR5 may be specific to this model, and not for angiotensin II for example. Further studies are necessary to dissect any difference amongst the hypertension models including 2-kidneys 1-clip and genetic models.

Aldosterone increases cardiac and renal CCR5^51, 54, 56^, but it is not clear if it does the same in arteries or endothelium and if CCR5 propagates any changes in the vasculature. Herein, we confirmed that aldosterone increases CCL5 and CCR5 and described CCL5 and CCR5 as novel regulators of endothelial homeostasis via adjusting Nox1 and NFκB in the vasculature. These findings corroborate our previous study^19^, where we demonstrated that CCL5, via CCR5, induces VSMC proliferation and migration via Nox1-derived ROS likely dependent on NFκB activation. The involvement of NADPH oxidases and NFκB activation in the genesis of aldosterone-induced cardiovascular injury has been described before^8, 11, 58^. NADPH oxidases are the major source of ROS in the vasculature and have been implicated in vascular dysfunction, inflammation, and remodeling by regulating NFκB, an important redox sensitive transcript factor, which regulates multiple inflammatory genes^59^. Furthermore, others have reported that NFκB can also regulate NADPH oxidase expression^40, 60^ indicating positive feedback between NADPH oxidases and NFκB.

While CCL5 can bind to CCR1, CCR3, CCR4 and CCR5, it has the highest affinity to CCR5^16^. By *in vitro* and *ex vivo* experiments, we examined whether CCL5 induces endothelial dysfunction and inflammation via CCR5. By treating arteries or endothelial cells with CCL5 with or without Maraviroc (CCR5 antagonist), we found that CCL5 impairs endothelial function and inflammation and increases NFkB activation and NOX1-derived ROS dependent on CCR5. Therefore, we can suggest that increases in circulating CCL5 might be activating CCR5 and triggering vascular injury and perhaps hypertension and renal damage in aldosterone treated mice.

It is important to acknowledge the limitations of the current study. First, we did not analyze immune cell infiltration into the arteries or kidney, which might be an additional mechanism whereby deficiency in CCR5 protects the vasculature and kidney. Instead, we mostly focused on a direct mechanism of CCL5 on endothelial cells. Second, we used aortae, and not resistance arteries. Blood pressure is normally regulated by small arteries, but multiple mouse models of hypertension are associated with vascular dysfunction in large^25, 61, 62^ and small vessels^8, 63, 64^, including aldosterone^8, 61, 62^, therefore we can suggest that vascular dysfunction might be displayed in small arteries too. Finally, we did not evaluate the source of CCL5. CCL5 is produced by many cell types including immune and vascular cells, therefore we cannot confirm that increases in CCL5 produced by aldosterone treatment is generated by an immune compartment or any other cell type. Future experiments including CCL5 deficiency in cell specific population or bone marrow transplant (CCL5 knockout mice into wild-type mice) could overcome this limitation.

Despite these limitations, our study provides the first evidence that aldosteronism impairs vascular function, induces vascular inflammation, and triggers hypertension and end-organ damage via regulating CCL5 and CCR5, which in turn activates NFkB and NOX1-derived ROS and diminishes nitric oxide formation. Taken together, these results indicate that blockage of CCR5 may confer protection against high levels of aldosterone-associated cardiovascular damage and hypertension.

### Perspective

Overall, our findings provide novel mechanistic insights into the underlying causes of vascular damage, hypertension, and end-organ injury in the pathophysiology of aldosteronism. In addition, our data present CCR5 blockage, e.g., Maraviroc, a drug broadly used in human immunodeficiency virus (HIV)^+^ patients^65^, as a possible new avenue for the treatment of cardiovascular dysfunction and hypertension associated with high levels of aldosterone such as aldosteronism, obesity, and hypertension.

**What Is New?**

- Increase in aldosterone levels, a key characteristic of primary aldosteronism, obesity, and hypertension, augments circulating CCL5 and vascular CCR5, whereas CCR5 deficiency protects from aldosterone-induced vascular injury, hypertension, and renal damage.
- CCL5/CCR5 triggers endothelial dysfunction and vascular inflammation via NFκB and NOX1 activation.

**What Is Relevant?**

- Our study has clinical implications by suggesting that CCR5 blockage can prevent the vascular and renal injuries and hypertension in diseases associated with high levels of aldosterone.

**Clinical/Pathophysiological Implications?**

- Maraviroc, a selective CCR5 antagonist approved by FDA for the treatment of HIV infection, may demonstrate clinical implications also to treat cardiovascular diseases specially associated with high levels of aldosterone.

## Acknowledgments

RM. Costa and T. Bruder-Nascimento participated in the conception, design of the work, acquisition of the data, analysis and interpretation of the data, and redaction of the manuscript. DM. Cerqueira, A. Bruder-Nascimento, JV. Alves, WAC. Awata, S. Singh, A. Kufner, E. Cifuentes-Pagano, participated in the acquisition and the interpretation of the data. PJ. Pagano and J. Ho participated in the design of the work and redaction of the manuscript.

## Sources of funding

This work was supported by The São Paulo Research Foundation (FAPESP, 2022/06639-2) to RMC, NHLBI-R00 (R00HL14013903) and startup funds from University of Pittsburgh to T.B.-N, and by NIDDK R01DK125015 to J.H.

## Disclosures

None

## Nonstandard Abbreviations and Acronyms

ACh: 4,5-Diaminofluorescein Diacetate (DAF-2 DA) Acetylcholine
CCL5: Chemokine (C-C motif) Ligand 5
DBP: Diastolic Blood Pressure
DMEM: Dulbecco’s Modified Eagle’s Medium
ELISA: Enzyme-Linked Immunosorbent Assay
LTA: Lotus Tetragonolobus Lectin
MAP: Mean arterial pressure
MR: Mineralocorticoid Receptors
NFκB: Nuclear Factor Kappa B
PBS-T: PBS containing 0.1% Tween-20
PE: Phenylephrine
PBS: Phosphate-Buffered Saline
ROS: Reactive Oxygen Species
RANTES: Regulated on Activation Normal T Cell Expressed and Secreted
SNP: Sodium Nitroprusside
SBP: Systolic Blood Pressure
T-Reg: T-Regulatory Cells
VSMC: Vascular Smooth Muscle Cells

## Notes

### Competing Interest Statement

The authors have declared no competing interest.

